# Identification and characterization of shifted G•U wobble pairs resulting from alternative protonation of RNA

**DOI:** 10.1101/2024.12.31.630957

**Authors:** Md. Sharear Saon, Catherine A. Douds, Andrew J. Veenis, Ashley N. Pearson, Neela H. Yennawar, Philip C. Bevilacqua

## Abstract

RNA can serve as an enzyme, small molecule sensor, and vaccine, and it may have been a conduit for the origin of life. Despite these profound functions, RNA is thought to have quite limited molecular diversity. A pressing question, therefore, is whether RNA can adopt novel molecular states that enhance its function. Covalent modifications of RNA have been demonstrated to augment biological function, but much less is known about non-covalent alterations such as novel protonated or tautomeric forms. Conventionally, a G•U wobble has the U shifted into the major groove. We used a cheminformatic approach to identify four structural families of shifted G•U wobbles in which the G instead resides in the major groove of RNA, which requires alternative tautomeric states of either base, or an anionic state of the U. We provide experimental support for these shifted G•U wobbles via the potent, and unconventional, *in vivo* reactivity of the U with dimethylsulfate (DMS) in three organisms. These shifted wobbles may play important functional roles and could serve as drug targets. Our cheminformatics approach is general and can be applied to identify alternative protonation states in other RNA motifs, as well as in DNA and proteins.

**Graphical Abstract:** 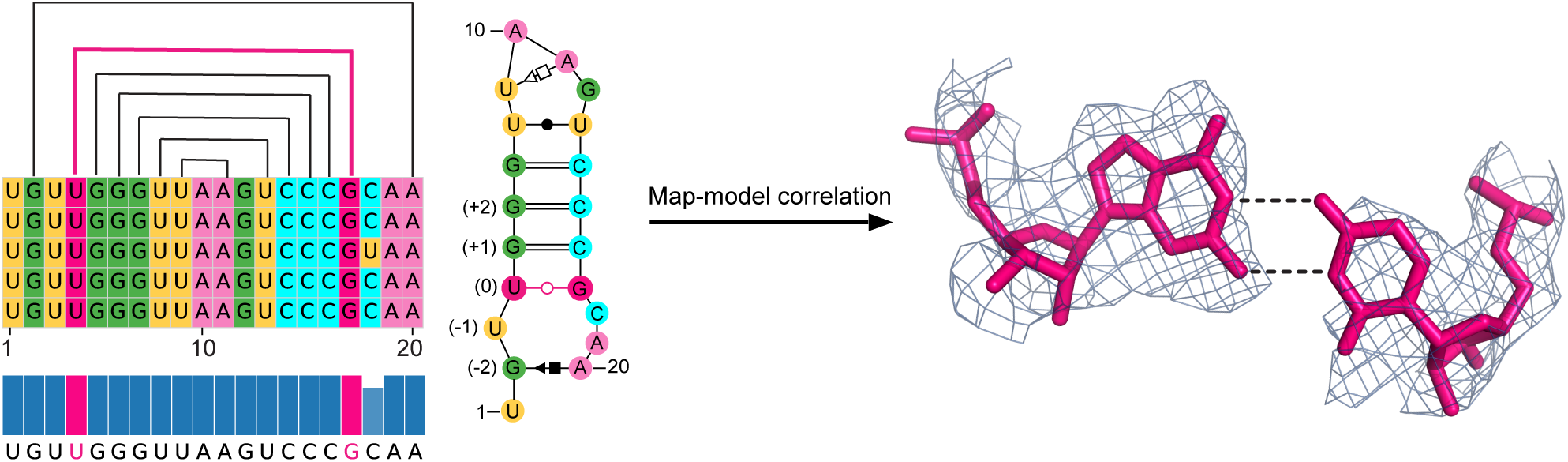

## Introduction

RNA is a relatively unassuming biopolymer comprised of only four similar sidechains and a simple sugar phosphate backbone. The nucleobase sidechains have only two sizes: the larger purines, A and G, which are comprised of fused five- and six-membered rings, and the smaller pyrimidines, C and U, which are comprised of a single six-membered ring. Additionally, the four nucleobases have highly similar functional groups of exocyclic keto and amino groups, and endocyclic imino nitrogens. Further limiting intermolecular interactions, the amino groups are aromatic, with their lone pairs are delocalized into the ring system. In many ways, this limited chemical diversity is incongruous with RNA’s prodigious functional diversity that includes catalysis (ribozymes), small molecule recognition (riboswitches), and synthesis of proteins (tRNAs, rRNAs, mRNAs) (1–3).

One way RNA is known to enhance its chemical range is through covalent modification. Indeed, tRNA and rRNA have long been known to have myriad covalent modifications such as base modifications (e.g. pseudouridine, 5-methyluridine, and 7-methylguanosine) as well as sugar modifications (e.g. 2′-O-methyl) (4–6). These modifications control numerous biological functions including codon recognition, reading frame maintenance, and tRNA decay. More recent studies have revealed extensive chemical modification of mRNAs as well. For instance, 6- methyladenosine, 8-oxo-guanosine, and pseudouridine have been found in mRNAs where they affect transcription, translation, and mRNA decay (7,8).

Less clear is whether non-covalent modifications have a role to play in enhancing the chemical diversity of RNA. The nucleobases have long been thought to resist ionization and tautomerization at neutral pH. For instance, the p*K*_a_ of the bases are removed from neutrality, being near 4 for the Watson-Crick-Franklin (WCF) faces of A and C and above 9 for those of G and U (9); moreover, these values shift further from neutrality upon formation of WCF base pairs owing to coupling with RNA folding (10,11). Watson and Crick reasoned that tautomers of the DNA bases must be rare in order to maintain the fidelity of base pairing (12), and calculations have indicated that tautomers of DNA and RNA are energetically unfavorable, with estimates of ∼5 to 10 kcal/mol penalties for the formation of the neutral tautomers of the bases (13–18). Nonetheless, there is experimental evidence, both direct and indirect, that ionization and tautomerization of the bases do indeed occur. Regarding ionization, we showed that C75 of the HDV ribozyme has a p*K*_a_ shifted up to 7.1 at biological concentrations Mg^2+^ (19,20), that C8 in the base quartet of beet western yellows virus has a p*K*_a_ of 8.1 (21), and that the A in an A^+^•C wobble pair can protonate at neutrality when the nearest-neighbors have strong WCF base pairing (22). Additionally, Lilley and colleagues showed that A1 in the twister ribozyme serves as the general acid to protonate the leaving group and has a pKa shifted towards 7 (23), and Wohnert and colleagues demonstrated that A11 in a quartet in a GTP-binding aptamer has a p*K*_a_ in the bound state of at least 8.9 (24). In all the above cases, the base is A or C and it forms a cationic state. We also provided indirect evidence that the bases and hydrated metal ions can ionize to perform chemistry in ribozymes (25,26). Regarding tautomerization, there has been indirect evidence of tautomers playing important roles in the mechanisms of the hairpin and HDV ribozymes (17,27,28) and direct evidence that the ribosome can form tautomers in a G•U pair (29). Ionization of the bases in so-called “reverse protonation” (26,30,31), as well as re-protonation of the bases in tautomerization, could lead to novel base-pairing, proton transfer in catalysis, and new sites for protein binding and specific drug targeting of RNA. Despite these anecdotal examples of ionization and tautomerization of the bases, there is little evidence as to whether it is widespread, present in anionic bases, or due to specific structures.

In this work we present a cheminformatic workflow that identifies and analyzes RNA structural motifs containing non-covalently modified residues. We focus on rare G•U wobbles (Figure 1) and do so in part because they are understudied but also because anionic bases, which can form in them, are rare in RNA. Recently, Cate and Westhof provided the first report that anionic GU pairs can form in bacterial rRNAs (32). Here, we significantly expand on this by employing a workflow to identify four separate structural clusters containing alternative forms of G•U wobbles across all three domains of life, in which the G, rather than the U, is shifted into the major groove. Secondary and 3D models are provided for each cluster, which reveal potential driving forces for formation of the shifted wobble including metal ion binding in the major groove and extensive intra and interstrand minor groove interactions within the local RNA fold as well as with amino acids. We also present dimethylsulfate (DMS) probing experiments in three organisms that support the WCF face of the U in a shifted G•U wobble as being deprotonated *in vivo*.

**Figure 1.**
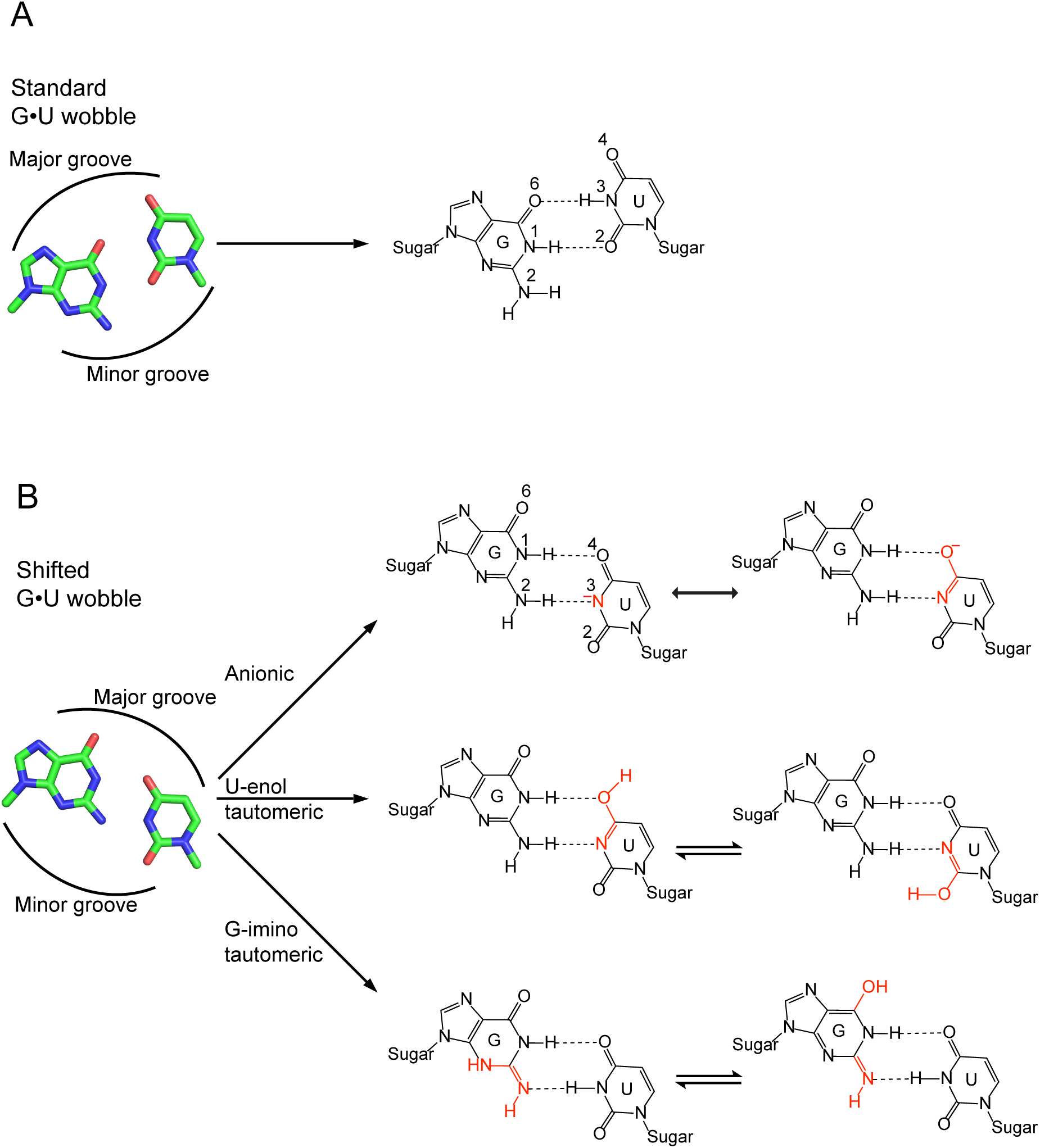
Structures of standard and shifted G•U wobbles. (A) Standard and (B) shifted wobbles. For the shifted wobbles, we provide charged (top) and tautomeric (middle and bottom) wobbles between G and U. For the charged wobble, we provide the enolate resonance form to the right. Positions of major and minor grooves are provided in both panels. For the stick drawings, the position of the sugar is depicted by a methyl group.

## Materials and Methods

### Workflow to identify non-redundant shifted G•U wobbles (From Figure 2A)

#### 1. Downloading and characterizing RNA structures

The workflow started with collecting structures containing RNA entities from the RCSB Protein Data Bank (PDB) (33). Structures were collected in the Crystallographic Information File (CIF) format. An initial resolution cut-off of 3.2 Å was applied to ensure satisfactory quality of structures. This resolution cut-off excluded all structures solved by solution NMR leaving just X-ray diffraction and cryo-EM structures. The selected structures were then characterized by Dissecting the Spatial Structure of RNA (DSSR) software (34). This step output base pair, hydrogen bond, stacking, glycosidic angle, and sugar pucker information for each structure file.

**Figure 2.**
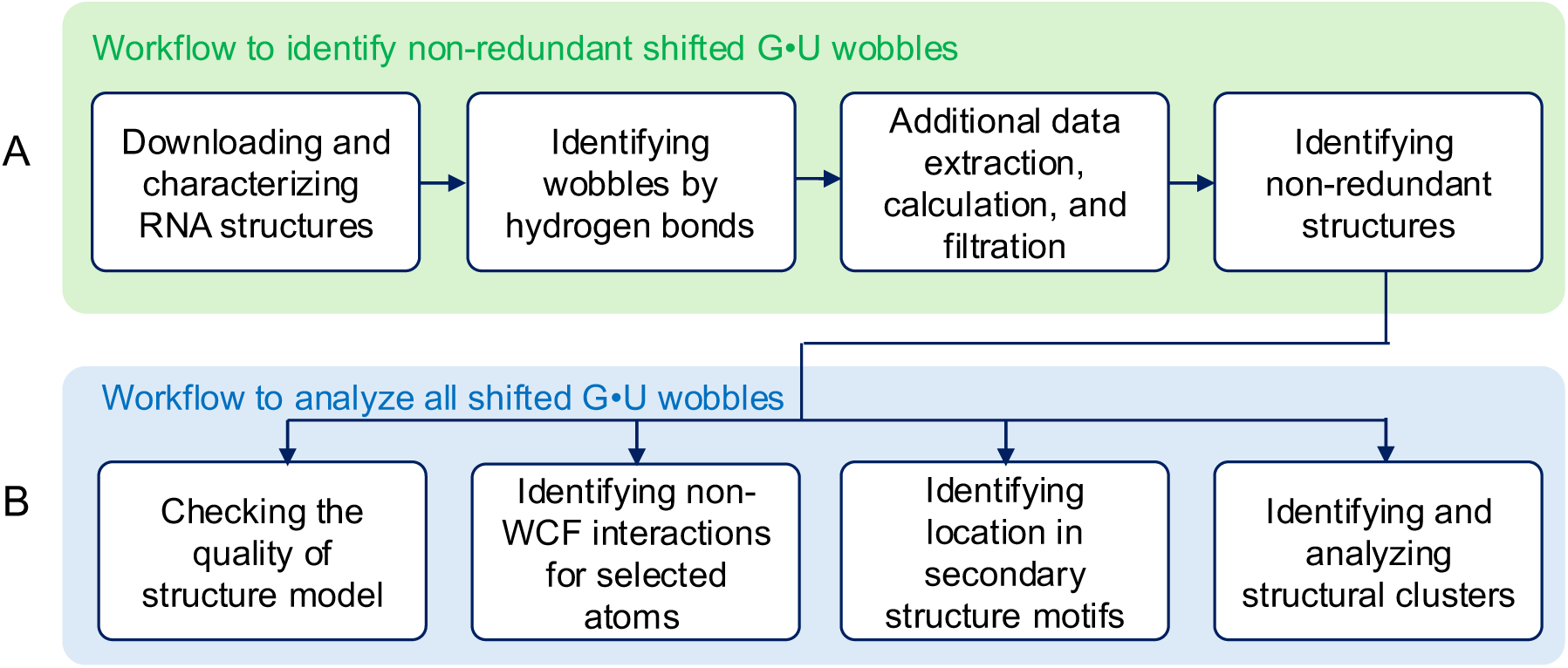
Workflow to identify and analyze shifted G•U wobbles. (A,B) Steps to (A) identify non-redundant shifted G•U wobbles and (B) analyze all shifted G•U wobbles.

#### 2. Identifying wobbles by hydrogen bonds

From the DSSR base pair information, all G•U base pairs were identified and filtered as wobble or non-wobble base pairs. All base pairs called by DSSR as G•U wobbles were considered for the next steps of the analysis as standard wobbles. Any base pairs containing hydrogen bonds between G(N1) and U(O4), as well as G(N2) and U(N3) (see Figure 1) were binned to shifted wobble base pairs.

#### 3. Additional data extraction, calculation, and filtration

For all the standard and shifted wobble base pairs, a two-step data extraction was performed. In the first step, metadata of the solved structures containing the wobbles were extracted from the RCSB PDB, including the name of source and expressed organisms, type of RNA molecule, segment ID, and length of the reference chain. In the next step, three types of information were calculated for each wobble: distances, dihedral angles, and average temperature factors. To calculate distances, three atoms from the WCF edge of G (O6, N1, and N2) and three atoms from the WCF edge of U (O4, N3, and O2) were considered. For each atom, the distances to the three WCF atoms of the other residue were calculated, resulting in a total of nine distances. To evaluate the coplanarity of the G and U, three dihedral angles were calculated: one along the O6-O4 distance and two along the distances of the hydrogen bonds required for each wobble. These latter two consist of the O6-N3 and N1-O2 distances for the standard wobbles and the N1-O4 and N2-N3 distances for the shifted wobbles. Finally, the average temperature factors for the nucleobase atoms of G and U forming the wobbles were calculated. We excluded cases where base pair forming residues were from different chains or where a residue index included an insertion code (i.e., a letter).

Using the calculated structural parameters, both standard and shifted wobble base pairs were evaluated for the quality of hydrogen bonds in solved structures through a three-step quality check process. Structures were required to have (1) hydrogen bonded distances of ≤ 3.4 Å and dihedral angles of ≤ 50°, (2) average temperature factors of ≤ 100 Å^2^ for G and U, (3) G(O6) to U(O4) distance of > 3.4 Å.

#### 4. Identifying non-redundant structures

From the dataset of structures, one typically finds a lack of uniformity of structures in terms of source organism, RNA type, and location of G•U wobbles. For instance, there are many ribosomal structures from some organisms (e.g., *Escherichia coli*) and no such structures from others. Therefore, for each group of structures from the same organism and RNA type with the same (or near identical) residue index, one representative structure was selected. Because the sequence indexes did not always match between structures of the same RNA in the same organism, indexes were first adjusted by alignment to one reference sequence for each organism and RNA type. Sequence alignments were performed with EMBOSS needle (35). Following are the steps to identify the representative structure for G•U wobbles from the same organism, RNA type, and adjusted residue indexes. The collection of representative structures constitutes our ‘non-redundant dataset’.

A. All groups contain one or more G•U wobble instances. For each of the wobble instances within a group, a six-residue motif (the G•U wobble plus one residue above and below both the G and U) was clipped from the originally solved structure. Instances of motifs with missing residues or atoms were excluded from further analysis if there are other instances within the same group that have all six residues and all atoms.
B. If there was only one instance of a G•U wobble within a group, then that motif was selected as the representative structure of the group. If there were only two instances, then the one with the lowest average temperature factors for G and U was selected as the representative structure. If there were more than two instances, then steps C and D were performed to select the representative structure.
C. The various instances of a motif within a group were compared with one another using pairwise RMSD values calculated using Biopython’s superimposer library (36). Because this calculation requires structures to contain the same number of atoms, we followed the recent approach by Kollmann and colleagues to coarse grain residues by selecting just five atoms from each residue: P, C4′, N9, C2, and C6 for purines and P, C4′, N1, C2, and C4 for pyrimidines (37).
D. From all instances within a group, one average structure was generated after aligning the structures of the instances. The instance with an RMSD that is closest to the average structure was selected as the representative structure.

### Workflow to analyze all shifted G•U wobble (From Figure 2B)

#### 1. Checking the quality of the structure model

Two additional quality assessments were performed: (1) Calculating the correlation coefficients between the experimental electron density maps and modeled structures and (2) Analyzing the hydrogen bond distances and angles of the two hydrogen bonds required for the shifted wobbles. These two quality checks were only performed for the shifted G•U wobbles of the non-redundant dataset.

For the first type of assessment, the electron density map and the modeled structure of the entire molecule for each non-redundant shifted G•U wobbles were obtained from the RCSB PDB (33). Using the Phenix software package, the map file was compared with the corresponding structure file to calculate map-model correlation coefficients (CC) (38). To assess the fit of the shifted G•U wobble to the experimental electron density map, the map-model CCs of the G and U residues were compared with the mean and median map-model CCs for all residues within the corresponding chain.

For the second type of assessment, hydrogen atoms were first added to a clipped structure only containing the shifted wobble using PyMOL. Then, the hydrogen-acceptor distances and the donor-hydrogen-acceptor angles for the G(N1)-U(O4) and G(N2)-U(N3) hydrogen bonds were calculated. We consider good hydrogen bonding geometries to exhibit distances of ≤ 2.5 Å and linear angles that are ≥ 140°.

#### 2. Identifying non-WCF interactions for selected atoms

Using PyMOL’s API along with python scripts, interactions within 3.4 Å of O2′, O4′, N3, N2, O6, and N7 of G and O2′, O4′, O2, and O4 of U were identified for all the non-redundant shifted G•U wobbles.

#### 3. Identifying location in secondary structure motifs

From the base pair information extracted from the DSSR characterization output, the non-redundant G•U wobbles were binned based on their location in one of the five secondary structure motifs: (1) inside stem, with one WCF base pair above and one below), (2) terminal, with at least one WCF base pair above, (3) terminal, with at least one WCF base pair below, (4) unstructured, where no WCF is right above or below and the wobble does not occur at the closing base pair of a hairpin with a maximum of 10 nucleotides, and (5) inside a loop.

#### 4. Identifying and analyzing structural clusters

To identify groups of non-redundant shifted G•U wobbles with similar structural orientation, the pairwise RMSDs of the corresponding six-residue motifs mentioned above were compared. The above coarse graining approach was followed to keep the number of atoms the same for the RMSD calculations. Hierarchical clustering was performed on the distance matrix using the centroid method, also known as Unweighted Pair Group Method with Arithmetic Mean, implemented in the SciPy library (39). To identify clusters of structures, an RMSD cut-off of 1.23 Å was identified on the basis of the similarity of the G and U residue indexes.

One consensus sequence and one secondary structure were assigned to each cluster. To this end, two different segments of the structure were prepared for each member of a cluster. The first segment includes three residues upstream from the G (or U), the G (or U) itself, the entire span of residues between the G and U (or U and G), the U (or G) itself, and three residues downstream from the U (or G). This structural segment was then characterized by DSSR (34) to extract the sequence, which was later used to perform multiple sequence alignment by pyMSAviz and assign a consensus sequence by following the Cavener rule (40). The second segment is the same as the first, except that the number of residues upstream from the G (or U) and downstream from the U (or G) may increase depending on the cluster and indicated in the Results. This segment was then characterized by DSSR to extract secondary structure and Leontis and Westhof notation (41). The most frequent secondary structures were assigned as the consensus secondary structure.

### Analysis of DMS-MaP probing data

Raw sequences were downloaded from the SRA using accessions provided in the Supplementary Table S1. Replicates with the same treatment condition were pooled, and reads were trimmed with cutadapt (42) according to the published methods for each library. Trimmed reads were then mapped and mutations were counted using the default settings of ShapeMapper2 (43). For *coli* and *S. cerevisiae*, raw reactivities were calculated by subtracting the mutation rate of the untreated sample from the mutation rate of the DMS-treated sample mutation rate. For *H. sapiens*, an untreated sample was not available, so the mutation rates of the DMS treated sample were used as raw reactivities. Finally, these raw reactivities were normalized by dividing each mutation rate by the average mutation rate of the 90^th^-98^th^ percentile. All reactivities over 1 (i.e. > average of the 90^th^-98^th^ percentile) were set to 1, all values under -0.1 were set to -0.1, and all nucleotides with raw background modification rates greater than 0.05% were discarded.

## Results

### Identifying shifted G•U wobble pairs

We were interested in identifying nucleobases with unusual protonation states (Figure 1). To do so we developed a cheminformatics approach and scanned the RCSB Protein Data Bank (PDB) (33) for RNAs with unconventional base pairing (Figure 2). As of April 2023, we downloaded 6,817 RNA structure files from the PDB in Crystallographic Information File (CIF) format (33). These structures were solved using several experimental methods: 3,729 by X-ray diffraction, 2,331 by cryogenic electron microscopy (cryo-EM), 737 by solution NMR, and 20 by other experimental techniques. In our cheminformatics approach, we first applied a 3.2 Å resolution cutoff, which removed approximately 42% of the structure files. The remaining 3,915 structure files, many of which were redundant (see below), included 3,038 by X-ray diffraction, 875 by cryo-EM, and 2 by fiber diffraction. We were curious about the distribution of base pairing type (i.e. AU, GC, and G•U) in these structures. To obtain this information, we turned to the software tool Dissecting the Spatial Structure of RNA (DSSR) (34), which takes RNA CIF files as inputs and provides a variety of structural features as outputs such as base pairing partner and type, hydrogen bond distances, and stacking interactions. We found that within these 3,915 structure files, there were 389,175 AU, 899,418 GC, and 159,047 G•U base pairs.

Next, we delved into the G•U base pairs. In the standard G•U wobble, the O4 of U is resident in the major groove (Figure 1A), but we also found shifted G•U wobbles where the O6 of G is instead resident in the major groove (Figure 1B). As depicted in Figure 1B, the shifted G•U wobble can be stabilized by either an anionic form of the U or a tautomeric form of either base. Because of our interest in bases with alternative protonation states, we analyzed our collection of G•U base pairs for shifted G•U wobbles. All 159,047 G•U pairs were analyzed for hydrogen bonds between G(N1) and U(O4), as well as between G(N2) and U(N3) (see Figure 1). Remarkably, this analysis resulted in the identification of 1,114 shifted G•U wobbles.

Next, we applied a stringent set of hydrogen bond distance, dihedral angle, and temperature factor cut-offs to both the standard and shifted G•U wobble pairs (see Materials and Methods), which resulted in 373 high confidence shifted G•U wobbles (Supplementary Table S3). Data on the standard G•U wobbles can be found in Supplementary Table S4. Within this dataset of high confidence shifted G•U wobbles, certain wobbles were found to be overrepresented (i.e. sharing the same organism, RNA type, and residue index). For instance, there were 137 examples of the shifted G•U wobble between U660 and G696 in the 16S rRNA of *Thermus thermophilus* (Table 1 and Supplementary Table S3). It is notable that there was just a single example of the standard G•U wobble between these same two residues (Table 1 and Supplementary Table S4), indicating a very strong propensity for formation of the shifted wobble. Notably, the vast overrepresentation of shifted G•U wobbles relative to standard G•U wobbles held for all cases with multiple examples of the shifted wobbles (Table 1).

**Table 1:**
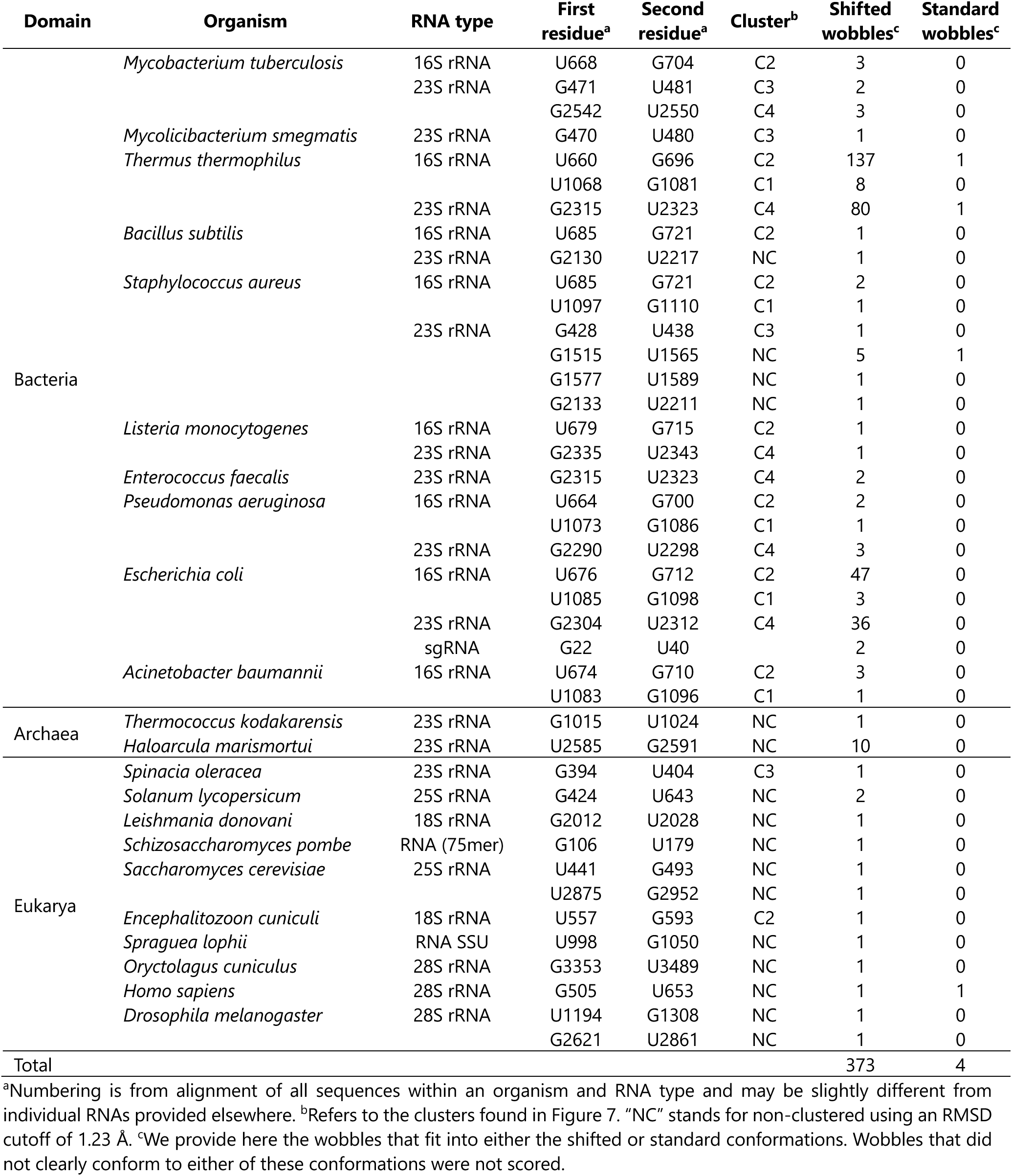
Frequency of shifted G•U wobbles in the full dataset.

While the overrepresented, high confidence shifted G•U wobbles were useful for assessing the prevalence of shifted over standard wobbles, they could lead to overcounting of unique shifted G•U wobbles. To address this, we developed a pipeline to identify a representative shifted G•U wobble for those examples sharing the same organism, RNA type, and residue index (Figure 2A). Upon filtering for redundancy, 41 unique examples of the shifted G•U wobble resulted (Table 1 and Supplementary Table S3). Among these, 27 were in bacterial RNAs from 10 species, 12 were in eukaryotic RNAs from 10 species, and 2 were in archaeal RNAs from 2 species (Figure 3 and Table 1). Regarding the distribution of these shifted G•U wobbles across RNA types, most were in rRNAs, with 23 examples in the large subunit and 16 in the small subunit.

**Figure 3.**
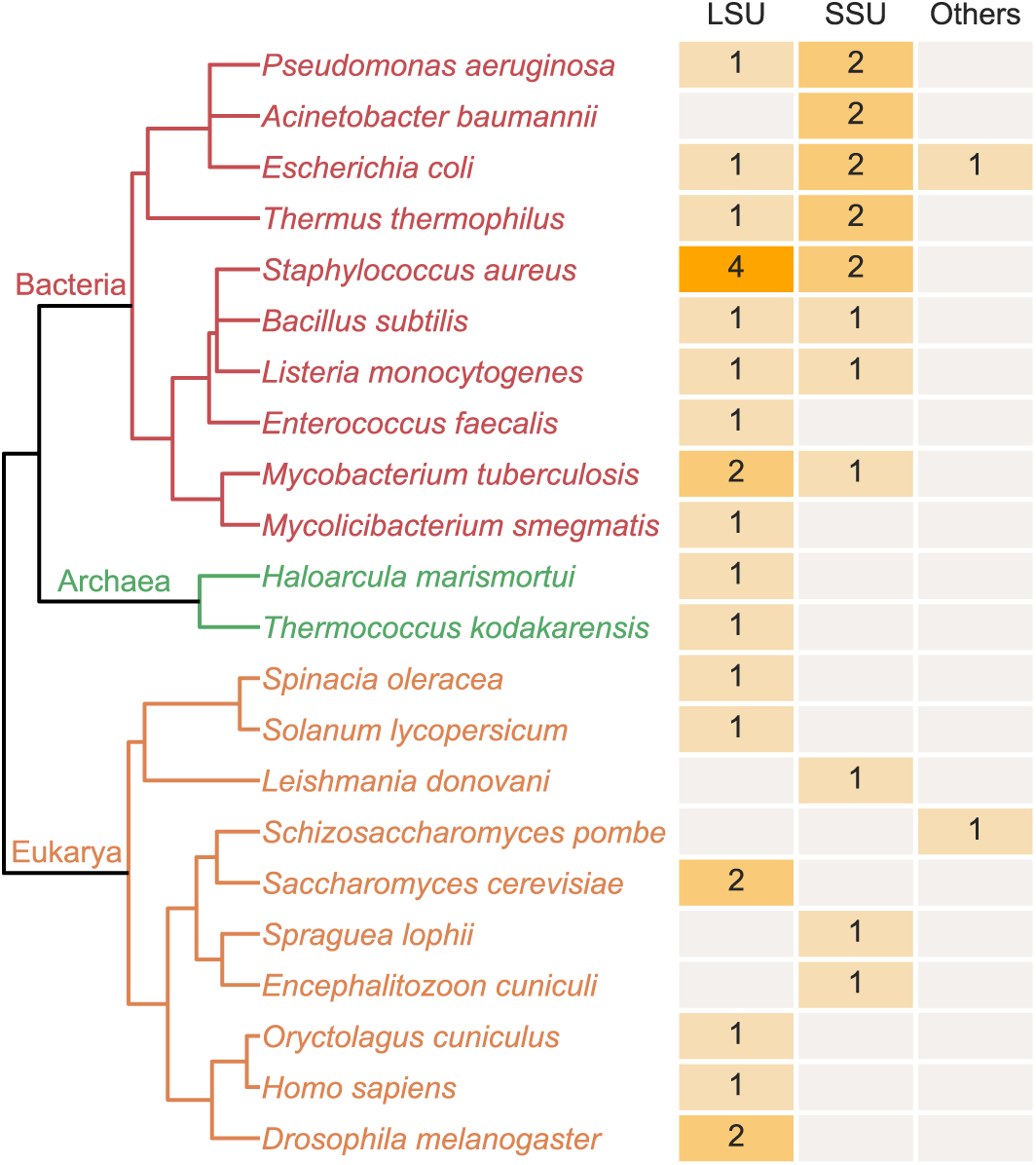
Distribution of shifted G•U wobbles across the three domains of life. Provided are RNA type and the source organism for the 41 non-redundant structures containing shifted G•U wobbles. ‘LSU’ indicates the large subunit of the ribosome and ‘SSU’ indicates the small subunit of the ribosome; ‘Others’ corresponds to a 75-mer and an sgRNA. The number of occurrences of each RNA type in each organism is provided with a heat map and a number. Gray boxes indicate that no data are available for the corresponding RNA types and organisms.

### Assessing the quality of the shifted G•U wobble pair assignments

The hydrogen bonds required for the formation of standard G•U wobbles are between G(O6) and U(N3), and between G(N1) and U(O2) (Figure 1A). In contrast, the hydrogen bonds necessary for the formation of the shifted G•U wobbles are between G(N1) and U(O4), and between G(N2) and U(N3) (Figure 1B). To judge the extent to which the standard and shifted G•U wobble candidates uniquely had these two sets of hydrogen bonds, we plotted the distribution of all non-redundant standard (n=6,636) and all non-redundant shifted (n=41) G•U wobbles for the nine possible pairwise distances between the three WCF face heteroatoms of G and three WCF face heteroatoms of U (Figure 4). The distance distributions for G(O6)-U(N3) and G(N1)-U(O2) were as expected for the standard and shifted wobbles, with small respective medians of 2.93 and 2.84 Å and narrow interquartile ranges (IQRs) of 0.20 and 0.22 Å for the standard G•U wobbles, but large respective medians of 5.57 and 5.68 Å and broad IQRs of 0.47 and 0.37 Å for the shifted G•U wobbles. Similarly, the distance distributions for G(N1)-U(O4) and G(N2)-U(N3) gave small respective medians of 3.02 and 3.20 Å and narrow IQRs of 0.33 and 0.31 Å for the shifted G•U wobbles, but large respective medians of 5.42 and 5.37 Å and broad IQRs of 0.33 and 0.37 Å for the standard G•U wobbles. This comparison of these two sets of distances between atoms participating in hydrogen bonds across the standard and shifted G•U wobbles enhances confidence in the assignment of the shifted wobbles. We note that the distances and IQRs are somewhat smaller and tighter, respectively, for the standard wobbles than the shifted ones, which may come from there being much more data for the standard wobbles (n_standard_=6,636 versus n_shifted_=41) or from researchers conducting model to electron density fitting refinements with standard protonation states. We also note that the medians from the other five pairwise combination of distances are above 3.4 Å line and so are not consistent with hydrogen bonds for either the standard or shifted G•U wobble pairs (Figure 4).

**Figure 4.**
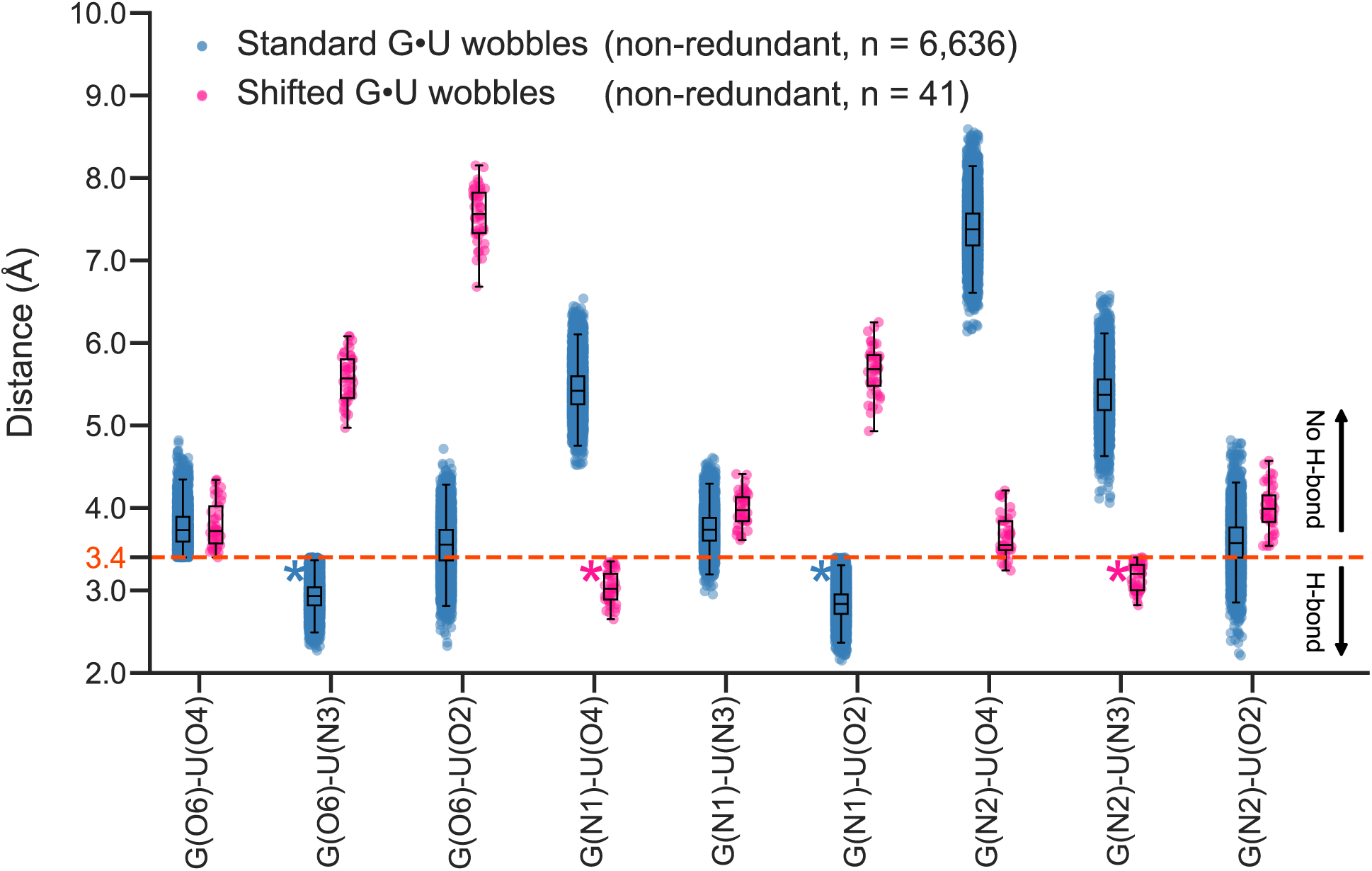
Distribution of distances between heteroatoms in all G•U wobbles. We provide distances for all nine possible combinations of the three WCF face heteroatoms of G (O6, N1, N2) and of U (O4, N3, O2). Combinations are provided in the order from major groove resident atoms to minor groove ones. The 6,636 non-redundant standard G•U wobbles are in blue and the 41 non-redundant shifted G•U wobbles are in red. The dashed orange line depicts the upper limit for a hydrogen bond of 3.4 Å. The standard G•U wobbles show two distinct distributions below this line, for G(O6)-U(N3) and G(N1)-U(O2) (blue asterisks), while the shifted G•U wobbles show two different distinct distributions below this line, for G(N1)-U(O4) and G(N2)- U(N3) (pink asterisks).

This lack of overlap between the distance distributions for the standard and shifted wobbles for the two sets of distances (Figure 4) supported the conclusion that the electron density map for shifted G•U wobbles cannot be accounted for by standard G•U wobbles. To further test this idea, we overlaid the shifted G•U molecular model on the electron density map for each of the 41 non-redundant examples of the shifted G•U wobble. An example is provided in Figure 5A, and all 41 examples are provided in Supplementary Figure S1.1-S1.41. In nearly all cases, the overlay of the model and the map was very good, supporting the interpretation that the shifted wobble is a unique molecular model to fit the electron density map. Next, for each of the 41 non-redundant examples, we plotted the raw map-model correlation coefficients as a function of residue index for all the residues in the structure, with an example provided in Figure 5B and all 41 examples provided in Supplementary Figure S1.1-S1.41. These plots revealed that the G and U of the shifted G•U wobble are generally in a well-defined region of the structure. Next, we plotted the distribution of each of the raw map-model correlation coefficients, with an example provided in Figure 5C and all 41 examples provided in Supplementary Figure S1.1-S1.41. Inspection of these examples confirmed that the G and U of the shifted G•U wobble have similar correlation coefficients as the rest of their structure. To visualize the correlation coefficient data for all 41 structures at once, we prepared box plot distribution of the correlation coefficients of the 82 G and U residues as well as of the 41 G residues alone and 41 U residues alone, both as raw and mean-normalized values (Figures 5D). The latter plots reveal that nearly all G and U residues have correlation coefficients similar to those of the rest of their structure. Three outlier G•Us were identified and are denoted in Supplementary Table S3. These were included in our analyses but did not make it into the clusters described below. In total, these data provide strong evidence that the electron density maps define a shifted G•U model with confidence.

**Figure 5.**
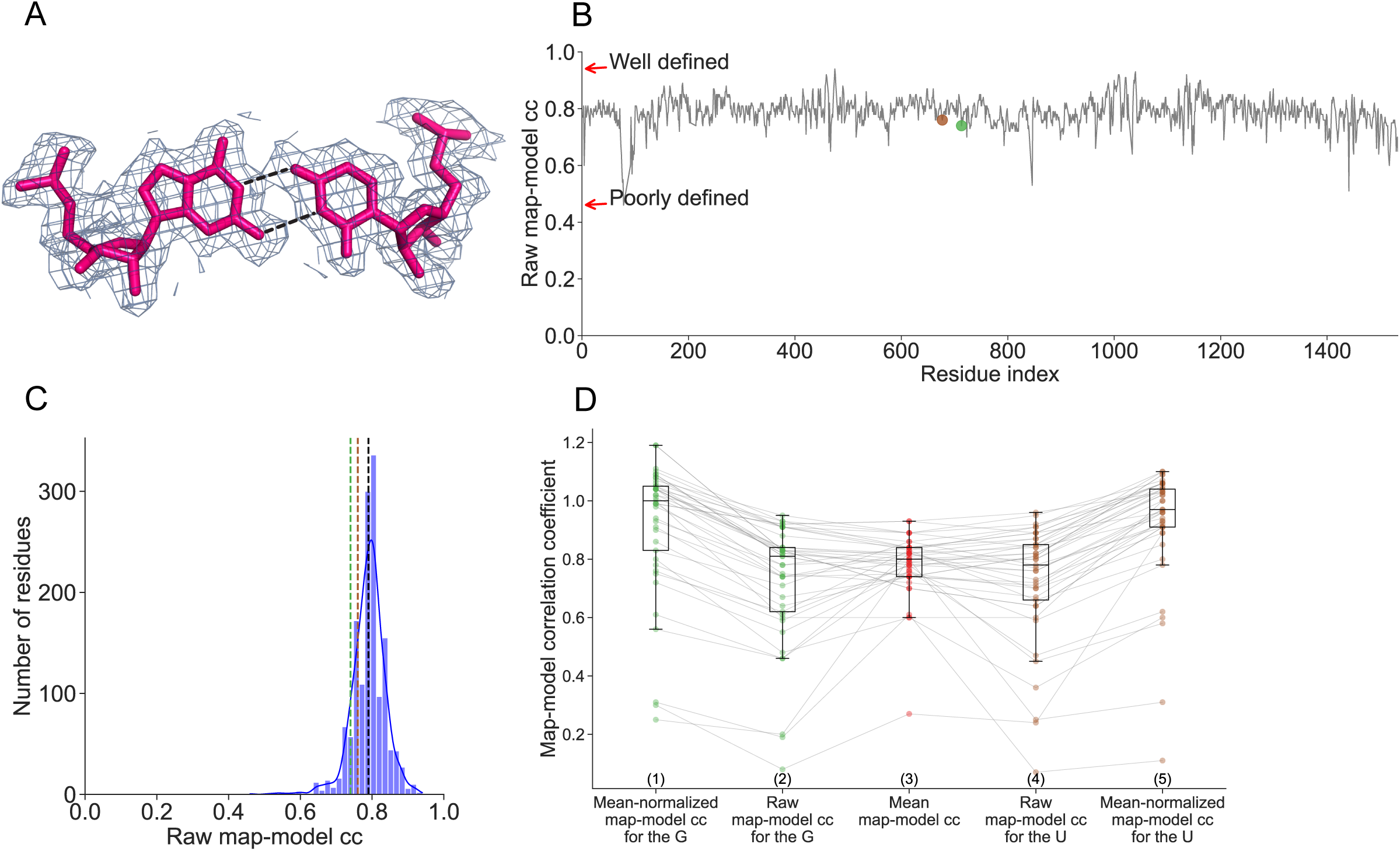
Map-to-model analyses for shifted G•U wobbles. (A) Sample model fitting to electron density map for 677U and 713G in 8B0X (chain ID: A). (B) Sample map-model correlation coefficient (cc) plot for the shifted G•U wobble from panel A. (C) Sample map-model cc distribution using the plot in panel (B). For panels B and C, the values for the G and U of the shifted wobble are denoted with green and umber dashed lines, respectively, and the overlapping mean and median are denoted with a single black dashed line. Panels (A)-(C) for each of the 41 non-redundant shifted G•U wobbles are provided in Supplementary Figure S1. (D) Distribution of map-model cc for all 41 shifted G•U wobbles. <u>Column 3:</u> Distribution of mean of map-model cc of all 82 nucleotides in the G•U wobble-containing chains. <u>Columns 2 and 4:</u> Distribution of raw map-model cc for the 41 G (Column 2) and the 41 U (Column 4) residues of the shifted wobble. <u>Columns 1 and 5:</u> Distribution of corrected map-model cc for the 41 G (Column 1) and the 41 U (Column 5) of the shifted wobble in which the raw value (Column 2 or 4) is divided by its mean value (Column 3). Gs are green and Us are umber.

Finally, to further assess the assignments of the 41 shifted G•U wobbles, we measured the distances and angles of its two hydrogen bonds upon the addition of a proton to the donating atom of each hydrogen bond. This approach has been shown to provide an additional evaluation of hydrogen bond quality (44). As shown in Supplementary Fig S2, 38 of these hydrogen bonds have high quality distances and angles, supporting their assignments as shifted G•U wobbles. Three outliers were again detected and are the same noted above.

### Sequences and secondary structures containing the shifted G•U wobble pairs

We were curious if there might be certain sequence and structural contexts that drive formation of the shifted wobble. To begin, we divided the 41 non-redundant shifted wobbles into G•U and U•G subclasses based upon which of the two residues, G or U, appeared first in the sequence (Figure 6A). We found that there was a near even split of the shifted wobbles between the two subclasses, with 22 members in the G•U subclass and 19 members in the U•G subclass (Figure 6B, C).

**Figure 6.**
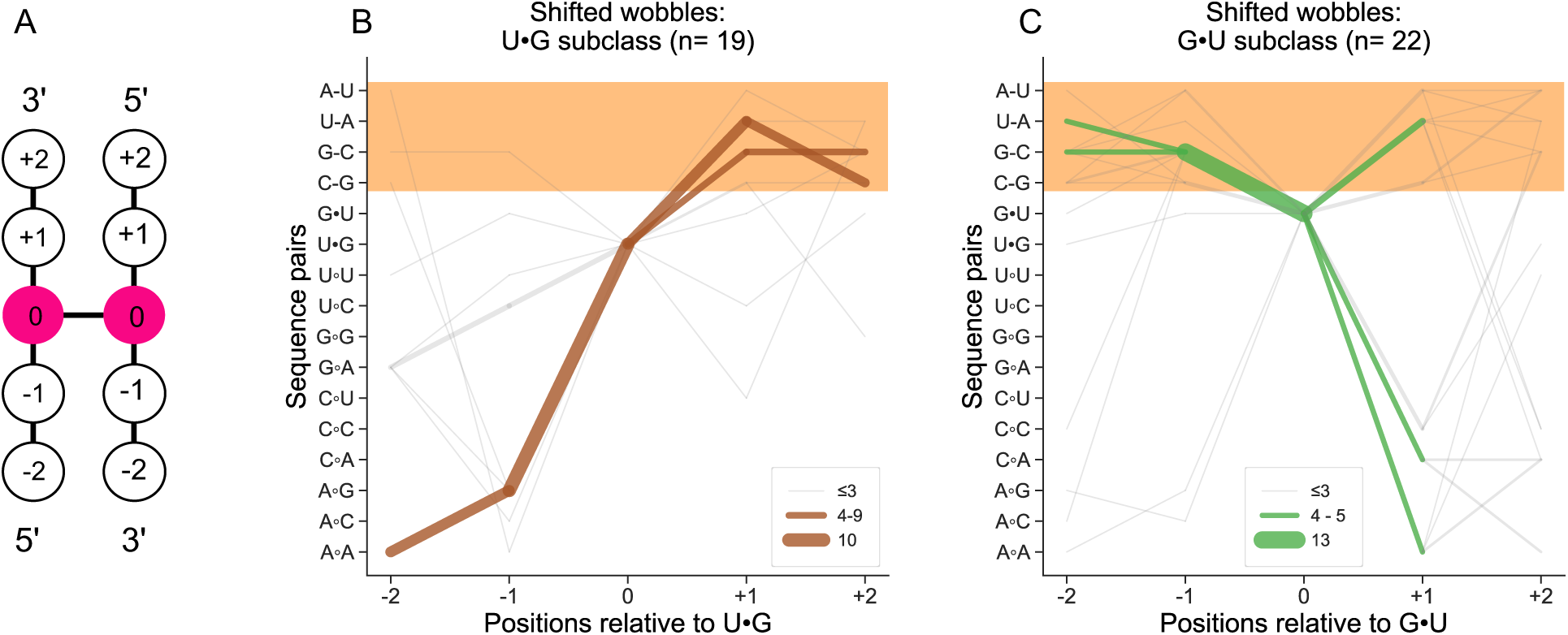
Patterns in sequences flanking the shifted wobbles. (A) Definition of positions of sequence pairs (defined as pairs of nucleobases opposite each other) relative to the shifted G•U wobble (pink circles) up to two steps above or below. The shifted wobble is defined as position 0, and positions below it are negative while those above it are positive. Panels (B) and (C) provide all 16 sequence pairs (y-axis) for all four positions flanking the shifted wobble (x-axis) for the G•U and U•G subclasses, respectively. The G•U and U•G subclasses were assigned on the basis of the first residue in the sequence. Line thickness scales with number of candidates and is consistent across the two panels. The orange field designates those sequence pairs with WCF complementarity, the majority of which form canonical WCF pairs (Supplementary Table S3).

We then inspected the sequences flanking the shifted wobble. The frequency of sequence pairs (defined as bases opposite each other but not necessarily base *paired*) for all 16 dinucleotide combinations are provided in Figure 6 for the four positions of -2, -1, +1, and +2, relative to the shifted wobble defined as position 0. We found that the vast majority—20 of the 22 G•U and 17 of the 19 U•G subclasses—contained at least one canonical WCF base pair as a nearest neighbor (Figure 6B, C, orange field.) For the 20 wobbles in the G•U subclass, canonical WCF base pairs were highly prevalent at both the -1 and +1 flanking positions, at 15 and 7 instances, respectively. Among these 20 structures, two contained canonical WCF base pairs on both sides of the shifted wobble. Turning to the 17 wobbles in the U•G subclass, canonical WCF base pairs were found predominantly at the +1 flanking position. In particular, the frequencies of canonical WCF base pairs at the -1 and +1 flanking positions were 1 and 16 instances, respectively. None of the U•G subclass structures had canonical WCF base pairs on both sides of the shifted wobble. The most frequent canonical WCF base pair at the -1 flanking position of the G•U subclass was GC, while UA was the most frequent canonical WCF base pair at the +1 flanking position of U•G subclass. The molecular basis behind these sequence and structural trends was investigated next.

### Clustering of the shifted G•U wobble pairs: Secondary structures, consensus sequences, and interactions

We were interested in whether the 41 non-redundant G•U wobble-containing sequences shared structural similarities. To address this, we constructed alignments of 3D structures, identified structural clusters, and converted these to consensus sequences and secondary structures. To start, we conducted an all-against-all structural alignment for these sequences (Supplementary Table S5) to sort them into clusters (Figure 7). Structural motifs used for clustering contained the -1, 0, and +1 sequence pairs as defined in Figure 6A and no other sequence. Briefly, we generated a distance tree by first coarse graining each of the six nucleotides in a structural motif and then applied hierarchical clustering to all 41 structures using the SciPy library (39) (see Materials and Methods). We then applied an RMSD cut-off of 1.23 Å, which resulted in four clusters having between 4 and 9 members each (Figure 7).

**Figure 7.**
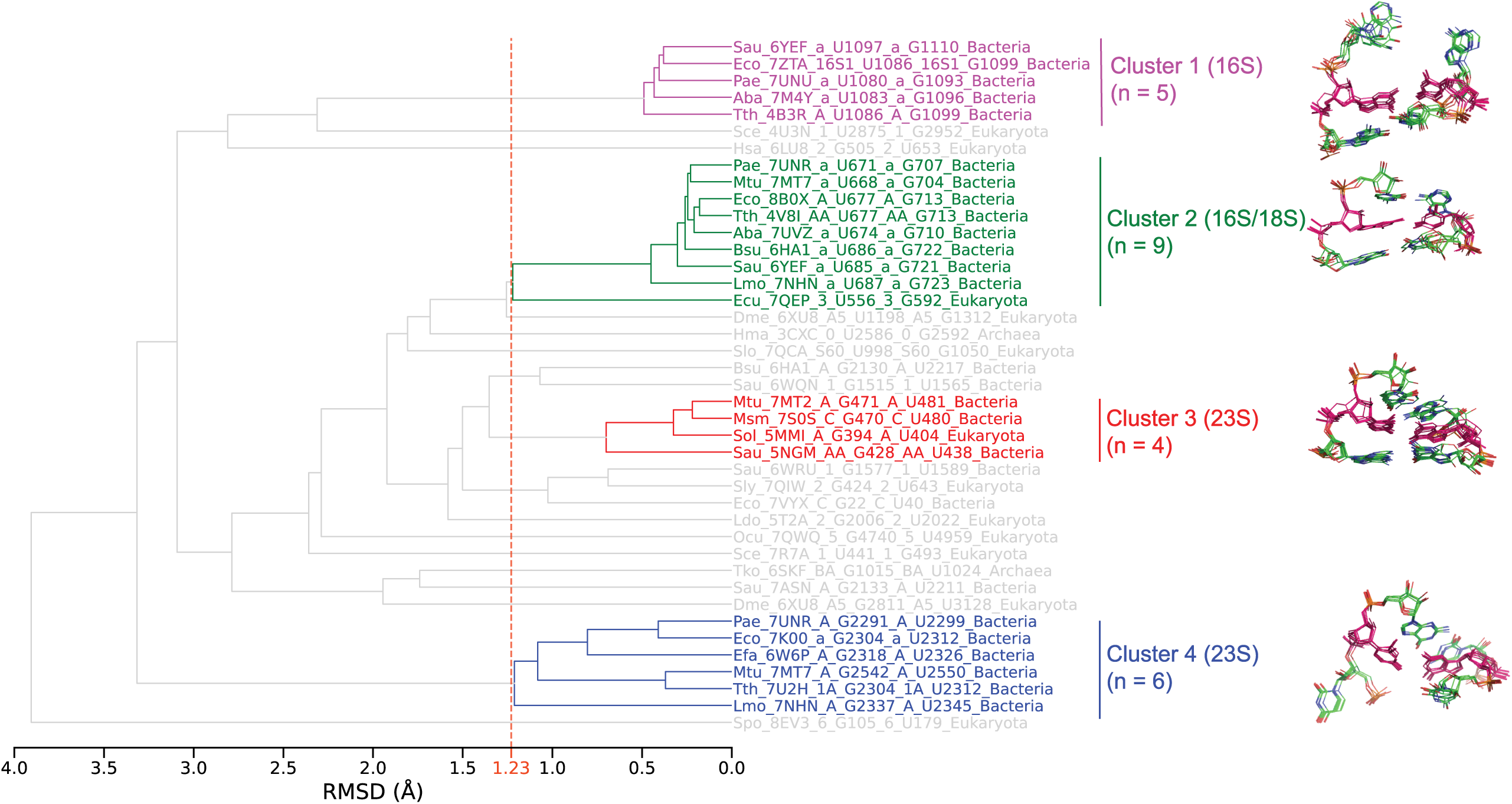
Distance tree showing the clustering of the non-redundant shifted G•U wobbles. For each of the 41 shifted wobbles (pink, right hand side), we included the -1 and +1 sequence pairs (green, right hand side) in the root mean square deviation (RMSD) calculation. Structural overlays (right hand side) show similarities within each cluster. The distances were calculated from the all-against-all structure alignment (see Materials and Methods). The vertical dashed orange line at 1.23 Å represents the upper limit in RMSD for sorting structures into the same cluster. Clusters were required to have at least four members, which resulted in clusters 1-4, with 5, 9, 4, and 6 members, respectively. The gray data remained unclustered at the 1.23 Å cutoff. The identity of each species is provided in Supplementary Table S3.

The first cluster was in the U•G subclass of shifted wobbles and had five closely related members from 16S rRNA bacterial species with an average distance between members of 0.45 Å (Figure 7, top). These were all at the equivalent position in 16S rRNA, with *E. coli* numbering of <u>U1086</u> and <u>G1099</u> (Figure 8A). (Note that we underline the G and U of the shifted wobbles to help with orientation.) Next, we conducted a multiple sequence alignment of these five members using pyMSAviz (see Materials and Methods). We included nucleotides starting at residue index -3 and ending at residue index +3 with respect to the shifted wobble position (Figure 8A). Thus for cluster 1, 20 residues from five sequences were aligned. This resulted in a near perfect alignment, with the only variation being at residue index +18 (Figure 8A, left). Next, for each of the five members, we retrieved the 3D structure of the 20 residues from the respective pdb files and obtained the underlying secondary structures for each of the five files in dot bracket notation using DSSR (34) (Figure 8A, right). In all five structures, the shifted U•G wobble was found at the base of a 4 base pair stem having three GC base pairs and enclosing a UUAAGU hexaloop. The three nucleotides before and after the shifted U•G wobble were largely unpaired in the crystal structures.

**Figure 8.**
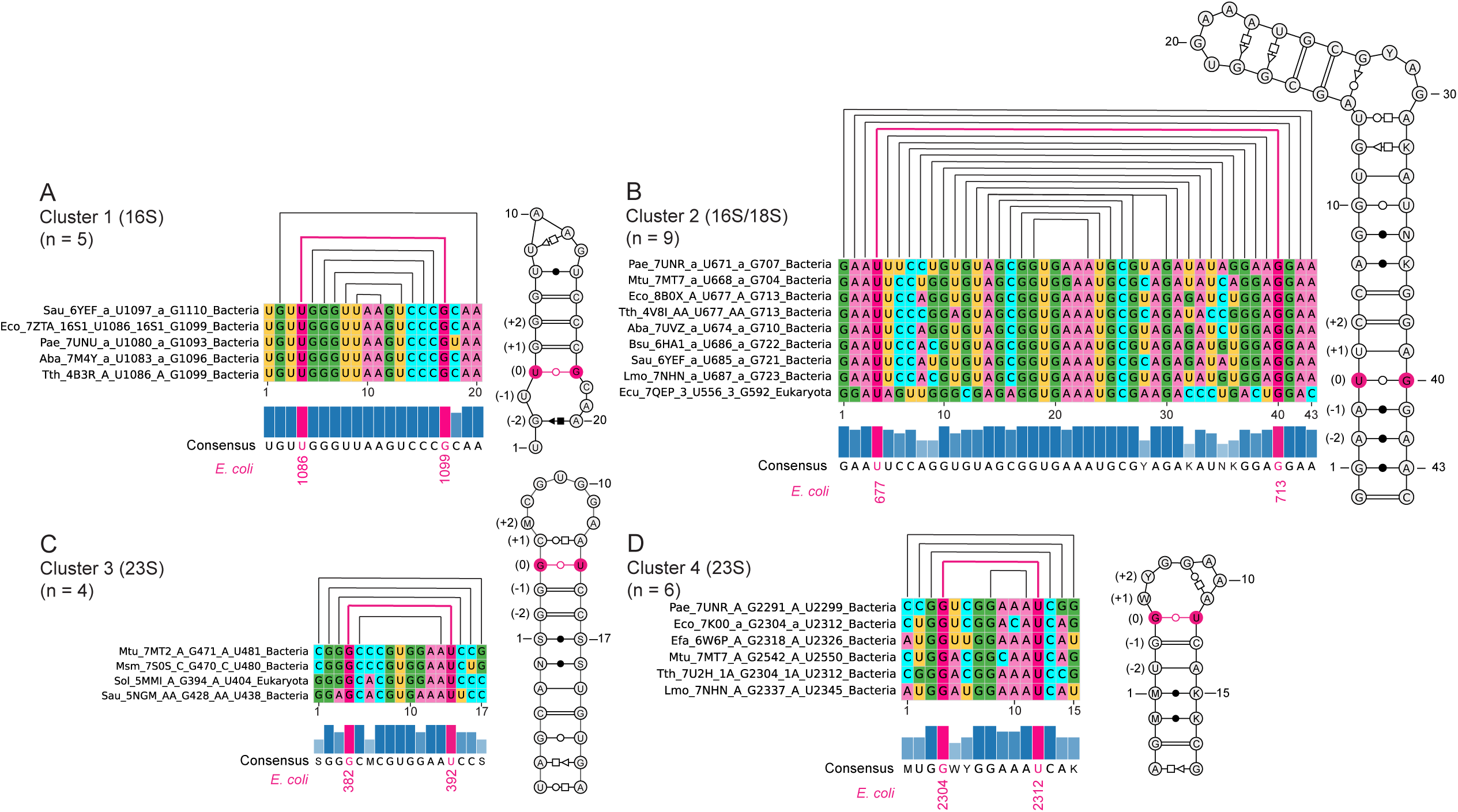
Consensus sequences and secondary structures for the four clusters for the non-redundant shifted G•U wobbles. Clusters 1-4 are provided in panels A-D, respectively. Sequence alignments, secondary structure representations, and consensus sequences were prepared as described in the Materials and Methods. The identity of each species is provided as a three-letter code to the left of each alignment and is also found in Supplementary Table S3. *E. coli* numbering for 16S and 23S rRNA, as appropriate, is provided in red below the shifted G and U in each alignment. Sequences were aligned from three residues before to three residues after the G•U. Position of sequence pairs within each cluster are provided in parentheses next to each secondary structure. Pairing is shown both above the alignment and in the secondary structure to the right of the alignment. As needed, stems were lengthened to accommodate additional base pairs found below the -3 sequence pairs, but these were not used in the alignment and so are not numbered. For the consensus sequence, the IUPAC code was used, where K= G or U; M= A or C; S= G or C; V= A, C, or G; W= A or U; and Y= C or U, and interactions of base pairs follow the convention of Leontis and Westhof (41).

To understand what interactions might drive formation of the shifted wobble in cluster 1, we analyzed the interactions of the heteroatoms between <u>U4</u> and <u>G17</u> using an interaction cutoff of 3.4 Å (Figure 9A). Briefly, this analysis considered pairwise interactions between an atom in the major groove, minor groove, and sugar atoms of the shifted wobble and an atom in the RNA chain, amino acids, metal ions, and water. We provide a stereoview of the minor groove and a stereoview of the major groove of each cluster in Supplementary Figure S3 to aid viewing. Beginning in the major groove of the shifted U•G wobble of cluster 1, the most frequent (4 of the 5 structures) interaction was a major groove-major groove intrastrand stacking interaction of between the oppositely charged <u>G17(O6)</u> and C16(N4). Also present in the major groove was an interaction (2 or 3 of the 5 structures) between the major groove atoms of <u>G17(O6)/U4(O4)</u> and a Mg^2+^ ion, which is also held in place by ligands from C18(N3) and the phosphate of U3. Note that we report the identity of the ion as modeled by the authors. The metal ion interaction suggests that this shifted G•U wobble might be in the anionic form (Figure 1) (see Discussion). In the minor groove, only one interaction was present (2 of 5 structures), which was a minor groove-minor groove interstrand stacking interaction between <u>G17(N3)</u> and G5(N1). Moving to the sugar region of the shifted U•G wobble, there was a hydrogen bond between <u>G17(O4′)</u> and C16(O2′). Looking at the “None” interaction column, it is notable that the <u>G17</u> base has many interactions, albeit not with its N7, while the <u>U4</u> base has none. We also see that the O6 of <u>G17</u> is always engaged in some interaction. Finally, we note that two of the five members in cluster 1 have multiple entries in the PDB. Specifically, 16S rRNA from *Escherichia coli* has 3 entries with a cluster 1 G•U shifted wobble and *Thermus thermophilus* has 8 entries with this shifted wobble, while neither has an entry with a standard wobble (Table 1 and Supplementary Table S3) supporting the shifted wobble. The second cluster identified was also in the U•G shifted wobble subclass and had nine members with an average distance of 0.51 Å between them (Figure 7, 2^nd^ set). Eight of the nine members were from bacterial 16S rRNA and were very closely related in structure, while the other member was a bit further related in structure and from eukaryotic 18S rRNA (PDB ID: 7QEP). All entries, including the eukaryotic one, were found at the equivalent position in 16S rRNA, with *E. coli* numbering of <u>U677</u> and <u>G713</u> (Figure 8B). The multiple sequence alignment of the 43 residues from these nine members, beginning three residues before the U•G and ending three after it, resulted in a strong consensus sequence, especially near the U•G wobble, of a UA pair above (8 of 9 sequences plus 1 AU pair) and an AG pair below (9 of 9 sequences) (Figure 8B, left). Again, we retrieved the 3D structures, here of 45 residues (one more residue from both ends to capture full length of the stem below the shifted U•G wobble) of all the nine members of cluster 2, and derived the secondary structures to identify the consensus secondary structure (Figure 8B, right). In all nine structures, the shifted U•G wobble was located within a 14 base pair stem that included a large (17 nt) structured loop. Cluster 2 has in common with cluster 1 a shifted U•G wobble at the base of a stem, but the two clusters differ in the nature of the bases above and below the wobble (Figure 8A and 8B).

**Figure 9.**
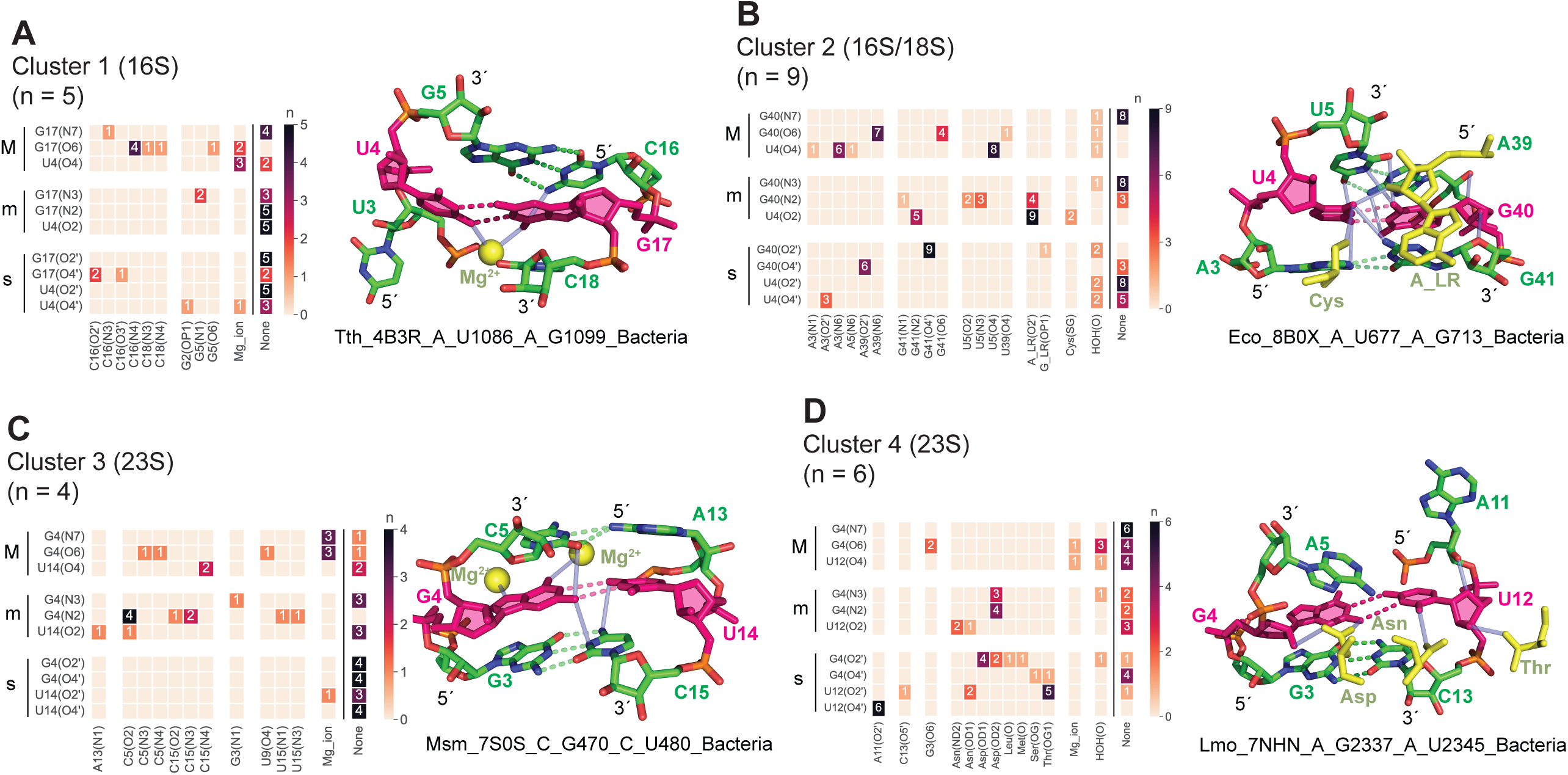
Interactions within each of the four clusters for the non-redundant shifted wobbles. Clusters 1-4 are provided in panels A-D, respectively. For each of the 41 shifted wobble (pink), we included the -1 and +1 sequence pairs (both in green), which were aligned as in Figure 6. In each panel, interactions diagrams and a representative structure are provided on the left and right, respectively; an overlay of all structures in a cluster is provided in Figure 7. For each interaction diagram, atoms from the shifted G and U are listed on the y-axis in the order: major groove (M), minor groove (m), sugar (s), while interacting atoms within 3.4 Å from the RNA chain, amino acids, metal ions, or water are provided on the x-axis, with the bases in alphabetical order nested by numerical order. For the representative structure, the base pairs are shown with dashed lines in the color of the respective bases, and vertical interactions are shown as solid gray lines. For clusters 1 and 3, we include metal ions that were frequent. For each structure, the view has the same orientation as in Figure 7, which leaves the minor groove in the foreground. Stereoviews of the minor and major groove of each structure are provided in Supplementary Figure S3.

Again, we considered the interactions of the shifted U•G wobble, here between <u>U4</u> and <u>G40</u> of cluster 2 (Figure 9B). First, we note that the UA above the U•G forms a standard WCF pair. Strikingly, the A and G of sequence pair -1 form a two-hydrogen bond cWW base pair (41), as do the AA and GA at sequence pairs -2 and -3, respectively (Figure 8B,9B). In the major groove of the shifted U•G wobble, frequent stacking interactions were observed with the adjacent base pairs. Specifically, the shifted U•G wobble stacks on the UA base pair above it: on the left strand, an intrastrand stacking interaction occurs between <u>U4(O4)</u> and U5(O4) in 8 out of 9 structures, while on the right strand, an intrastrand stacking was identified between <u>G40(O6)</u> and A39(N6) in 7 out of 9 structures. Additionally, extensive stacking interactions were observed with the AG base pair below the wobble: on the left strand, stacking was noted between <u>U4(O4)</u> and A3(N6) in 6 out of 9 structures, and on the right strand, between <u>G40(O6)</u> and G41(O6) in 4 out of 9 structures. These intrastrand stacking interactions with residues both above and below the shifted wobble contribute significantly to stabilizing the structure. In the minor groove, there was one dominant interaction (9 of 9 structures) of a hydrogen bond involving <u>U4(O2)</u> and A_LR_(O2′), where “LR” stands for long-range, which is for any residues outside the aligned region. The <u>U4(O2)</u> also interacted with G41(N2) (5 of 9 structures). These <u>U4(O2)</u> interactions are likely important for holding the U of the U•G wobble in the minor groove (see Discussion). In the sugar region, there were two dominant interactions: one was a hydrogen bond between <u>G40(O2′)</u> and G41(O4′) (9 of 9 structures), and the other is a hydrogen bond between <u>G40(O4′)</u> and A39(O2′) (6 of 9 structures). There was also a sugar-sugar interaction of <u>U4(O4′)</u> and A3(O2′) (3 of 9 structures). This extensive intrastrand involvement of the O4′-to-O2′ sugar hydrogen bonds, especially the O2′ and O4′ of <u>G40</u>, which interacts with the O4′ and O2′, respectively, of sugars below and above the shifted wobble along both strands, is particularly notable. This network of ribose interactions is like a ribose zipper, which conventionally holds together two *different* strands together using 2′OH interstrand interactions (45) but differs in that it also involves the O4′ and in that all the interactions are intrastrand. Looking at the “None” column, it is notable that <u>G40(N7)</u> of the shifted wobble again has no interactions, while <u>G40(O6)</u> is still strongly engaged. Unique to cluster 2 is the heavy engagement of the U of the shifted wobble in interactions, including its major groove resident O4 and minor groove resident O2. Notably, there is no high occupancy metal ion present in these structures. Finally, we note that there are a number of other cluster 2 16S rRNA entries in the full dataset (Table 1 and Supplementary Table S3). These include *E. coli* (47 entries) and *T. thermophilus* (137 entries); for the other members in this cluster, there were three or fewer structures available for each in the redundant dataset. Out of nine members in this cluster having nearly 200 shifted G•U wobble entries, there was only one entry forming a standard G•U wobble (Table 1), strongly supporting the shifted wobble.

Whereas the first two identified clusters were from the shifted U•G subclass of wobbles, the third cluster was from the shifted G•U subclass of wobbles, which had four members with an average distance of 0.50 Å between them (Figure 7, 3^rd^ set). These members were all from 23S rRNA rather than 16S rRNA, with three of the four being from bacteria and the other from Eukarya (PDB ID: 5MMI). All entries, including the eukaryotic one, were found at the equivalent position in 23S rRNA, with *E. coli* numbering of <u>G382</u> and <u>U392</u> (Figure 8C). The multiple sequence alignment of all 17 residues from the four members of cluster 3, again beginning 3 residues before the G•U wobble and ending 3 after it, resulted in a strong consensus sequence near the G•U of a CA pair above (4 of 4 sequences) and a GC pair below (3 of 4 sequences and 1 AU) (Figure 8C, left). We then retrieved the 3D structures of the 29 residues (six more residues from both ends to capture the full length of the stem below the shifted G•U wobble) of all the four members of cluster 3 and derived the secondary structures, with consensus stem-loop secondary structure found in Figure 8C, right. In three out of four structures, the shifted G•U wobble was found within a stem of 11 base pairs where there is one base pair above and nine base pairs below the shifted G•U wobble, and the wobble closed a loop of 9 nt with the sequence CMCGUGGAA (M= A or C).

Next, we considered the interactions of the shifted G•U wobble between <u>G4</u> and <u>U14</u> of cluster 3 (Figure 9C). First, we note that the GC of sequence pair -1 from below the G•U forms a standard WCF pair. Interestingly, the C and A of sequence pair +1 from above the G•U forms a two-hydrogen bond trans-WCF-Hoogsteen (tWH) pair (41) in which A13(N6) donates one of its protons to the C5(N3) and the other to the C5(O2), which differs from the more conventional AC wobble in which A(N1) has to be protonated (22). In the major groove of the shifted G•U wobble, there were limited RNA-RNA interactions, with only major groove-major groove intrastrand stacking between complementary charges on the adjacent <u>U14(O4)</u> and C15(N4) (2 of 4 structures) having more than one occurrence. However, like cluster 1, the major groove did host Mg^2+^, in this case two metal ions, with one metal ion interacting with <u>G4(O6)</u> and the other metal ion interacting with <u>G4(N7)</u>. This suggests that, like in cluster 1, this shifted G•U wobble might be in the anionic form (see Discussion). In the minor groove, there was only one dominant RNA-RNA interaction (4 of 4 structures), which involved a minor groove-minor groove intrastrand stack of complementary charges on <u>G4(N2)</u> and C5(O2). Notably, this minor groove intrastrand stack was in the left strand, while the major groove intrastrand stack was in the right strand, providing support for both strands of the structure. In the sugar region, there were no notable interactions. It can be noted that cluster 3 is like cluster 1 in that both are dominated by major groove interactions with a metal ion and both have relatively few minor groove and sugar interactions. Indeed rotation of cluster 1 about its axis coming out of the page reveals similarities, although the lower base pair is a GC in cluster 3 but a CG in cluster 1. Finally, there were not many additional cluster 3 23S rRNA entries in the full dataset, except for the shifted wobble in 23S rRNA of *Mycobacterium tuberculosis*, which has two structures in the redundant structures (Table 1 and Supplementary Table S3). Nonetheless, there were no examples of a standard G•U wobble by the corresponding residues, supporting the shifted wobble (Table 1).

The fourth and final cluster was also in the G•U shifted wobble subclass, where it had six members, which were related by an average distance of 1.0 Å (Figure 7, bottom). The six members were all from bacteria and were found at the equivalent position in 23S rRNA, with *E. coli* numbering of <u>G2304</u> and <u>U2312</u> (Figure 8D). The multiple sequence alignment of all 15 residues from the six members of cluster 4, starting 3 residues before the GU wobble and ending 3 residues after it, resulted in a strong consensus sequence near the G•U consisting of a WA pair above and a GC pair below (4 of 4 sequences) the G•U (Figure 8D, left). Upon retrieving the 3D structures of the 21 residues (three more residues from both ends to capture the full length of the stem below the shifted G•U wobble) of all the six members of cluster 4 and deriving the secondary structures, we ended up with a consensus stem-loop (Figure 8D, right). In all cases, the shifted G•U wobble was found atop a stem having six base pairs below, with a standard GC pair neighboring the G•U, and a 7 nt loop with the sequence WYGGAAA (W= A or U, and Y= C or U).

Next, we investigated the interactions of the shifted G•U wobble between <u>G4</u> and <u>U12</u> of cluster 4 (Figure 9D). As with the shifted G•U wobble from cluster 3, the GC of sequence pair -1 forms a standard WCF pair, but unlike cluster 3, there is no base pairing above the wobble. Very few interactions occur in the major groove. The only one of some note is a major groove-major groove intrastrand stacking interaction between <u>G4</u>(O6) and G3(O6) (2 of 6 structures). A metal ion was found with major groove atoms of the wobble, <u>G4</u>(O6) and <u>U12</u>(O4), but it was present in only one structure and so is not shown. The minor groove has more interactions and in this sense is somewhat reminiscent of cluster 2; moreover, in both clusters 2 and 4 most of the minor groove interaction are to species distal in sequence. In cluster 2, the dominant minor groove interaction was a hydrogen bond with the O2′ of a long-range A, while in cluster 4 all the minor groove interactions are to amino acids from UL5, a universally conserved protein from the large subunit (46). In particular, there is an aspartate that interacts with the N2 (4 of 7 structures) and N3 (3 of 7 structures) of <u>G4</u>, as well as an asparagine that interacts with <u>U12</u>(O2). In the sugar region, there is a multitude of interactions, again reminiscent of cluster 2. The dominant interaction (6 of 6 structures) is a hydrogen bond between <u>U12</u>(O4′) and A11(O2′) that is akin the hydrogen bond between <u>U4</u>(O4′) and A3(O2′) in cluster 2. The other interactions in the sugar region are again with amino acids, with the two dominant interactions involving <u>G4</u>(O2′) and the same aspartate that interacted with N2 and N3 of <u>G4</u>, and U12(O2′) and a threonine. Notably, <u>U12</u> engages its O2′ and O4′ in 5 of 6 and 6 of 6 possible times, respectively, presumably helping to anchor the U of the wobble in the minor groove as is characteristic of the shifted G•U wobble (see Discussion). We note that there are multiple other cluster 4 23S rRNA entries in the full dataset (Table 1 and Supplementary Table S3). In particular, there are 36 entries from *E. coli* and 80 from *T. thermophilus*. For the other members in this cluster, there were three or fewer structures available for each in the redundant dataset. For the more than 100 members in this cluster, there was only one member forming the standard G•U wobble (Table 1), strongly supporting the shifted wobble.

### Experimental support for the shifted G•U wobble pairs

In an effort to test whether the predicted G•U wobble pairs identified herein by cheminformatics are deprotonated in solution, we examined the dimethylsulfate (DMS) reactivity of ribosomes from three different organisms treated *in vivo* with DMS and read out using mutational profiling. Typically, DMS reacts with the WCF of A and C, which are deprotonated and can act as nucleophiles to attack a methyl group on DMS (47,48). Conventionally DMS should not react with G or U, as the imino nitrogen atoms on their WCF face are protected by protonation. However, if the U adopts either the anionic or enolic alternative protonation form (Figure 1B), it becomes deprotonated on the WCF face and has the potential to attack DMS, as confirmed by *in vitro* DMS experiments at elevated pH (49). We note that reactivity with a shifted G•U wobble may necessitate opening of the base pair, followed by rapid reaction with DMS before it closes back up.

Here we analyze published rRNA datasets for the reactivity of DMS with G and U in *E. coli* (50) and in *S. cerevisiae* and *H. sapiens* (51). Our analysis revealed significant DMS reactivity for three out of the six shifted G•U wobbles with available reactivity data; in all three cases, it was the U of the shifted G•U wobble that reacted (Supplementary Figure S4). In *E. coli*, we identified three shifted G•U wobbles through our cheminformatic approach (three rows in Table 1), and one of these was highly reactive with DMS *in vivo* (Figure 10 and Supplementary Figure S4). This wobble, consisting of U677 and G713 in 16S rRNA, was identified as being in Cluster 2 (Figures 7 and 8B). Supplementary Figure S4 shows its reactivity in the context of all U and G reactivities in *E. coli* rRNA. When compared to average DMS reactivities for all Us, U677 had higher DMS reactivity with a p-value < 0.0005 (Table S2). Reactivities of other Us and Gs in shifted wobbles identified in *E. coli* were insignificantly increased over the average (Supplementary Figure S4A). Nonetheless, although not significant, U1086 and G1099 in 16S rRNA of *E. coli* had moderate reactivity with DMS (Supplementary Figure S4A and S5A) suggestive of a U enol tautomer, possibly with deprotonation of G(N1) by U(O4), which would itself be externally deprotonated (Figure 1). Overall, the significant DMS reactivity of <u>U677</u> of a shifted G•U wobble strongly supports this U being either anionic or enolic (Figure 1B).

**Figure 10.**
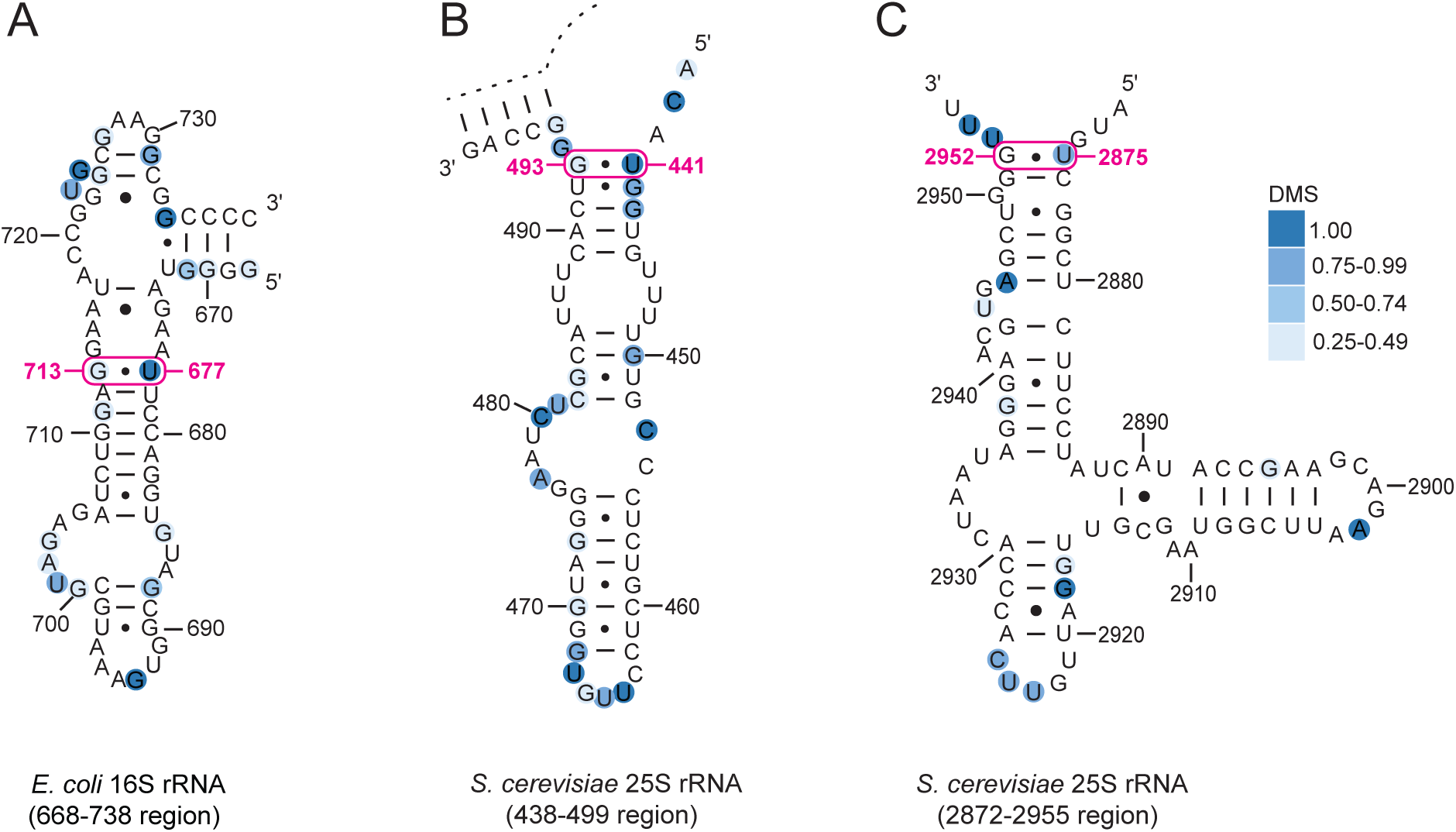
*In vivo* reactivity of DMS-treated rRNAs. (A-C) Secondary structures illustrating the reactivity of DMS in the (A) 668-738 region of *E. coli* 16 S rRNA, highlighting the high reactivity of U677 in a shifted G•U wobble, (B) 438-499 region of *S. cerevisiae* 25S rRNA, highlighting the high reactivity of U441 in a shifted G•U wobble, and (C) 2872-2955 region of *S. cerevisiae* 25S rRNA, highlighting the high reactivity of U2875 in a shifted G•U wobble. Additional examples are found in Supplementary Figure S5.

Next, we assessed the *in vivo* DMS reactivity of shifted G•U wobbles in the other two organisms, yeast and human. We identified two shifted G•U wobbles in *S. cerevisiae* 25S rRNA (two rows in Table 1). Strikingly, both had significant reactivity of the U (Figure 10B,C), with p-values of 0.0015 for <u>U441</u>•<u>G493</u> and 0.018 <u>U2875</u>•<u>G2952</u> (Figure S4, Supplementary Table S2). Like with <u>U677</u>•<u>G713</u> in *E. coli*, both <u>U441</u>•<u>G493</u> and <u>U2875</u>•<u>G2952</u> are terminal U•G wobbles. However, unlike <u>U677</u>•<u>G713</u> of *E. coli*, neither of these pairs in *S. cerevisiae* are part of a structural cluster (Figure 7), and the base pair between <u>U2875</u> and <u>G2952</u> by crystal structures is not reported by covariance analysis (52). Thus, while the high reactivity and consistent structure within bacterial ribosomes of *E. coli* U677•G713 suggests the deprotonated U is a conserved structural feature of bacterial ribosomes, the same is unclear for eukaryotes, although this may be due to the dearth of eukaryotic ribosomes represented in the protein databank. Finally, the shifted G•U wobble in humans between G505 and U653 did not show significant reactivity with DMS (Figure S4, S5).

In summary, there was significant *in vivo* DMS reactivity for three of the six shifted G•U wobbles identified herein that have available genome-wide DMS chemical probing datasets, and the reactivity was always with the U of the G•U wobble. This observation supports either the anionic U or enolic U forms of the tautomer (Figure 1). The three shifted G•U wobbles that did not react significantly with DMS may have more limited opening of the base pair. Overall, these data provide *in vivo* experimental support for the shifted G•U wobbles identified herein by computational means.

## Discussion

In this study, we provided a new approach for discovering RNAs with unconventional protonation and tautomeric states. Briefly, our approach is to draw the Lewis structure of a base pair intentionally having hydrogen bonding clashes, with the notion that the clashes can be resolved either by ionizing a base or by shifting a proton from one atom on a base to another atom on the same base as a tautomer, effectively swapping hydrogen bond donating and accepting roles. These changes can resolve one or even two hydrogen bonding clashes, for example if there are adjacent DD and AA clashes, where “D” is hydrogen bond donor and “A” is hydrogen bond acceptor. We then conduct 3D searches for the heteroatoms of such novel base pairs in the PDB, which is rife with base pairs; for instance, herein, we examined nearly 160,000 GU pairs. We then analyze the shifted wobbles to assure that they have good map-to-model coefficients both internally and relative to the rest of the structure, as well as favorable hydrogen bonding distances and angles, and extract their structural features. We then assign them to secondary structure motifs and identify structural clusters. In so doing we identified novel G•U pairs that require alternative protonations.

The shifted G•U pairs identified herein were found in all three domains of life, including multiple examples from bacteria and eukarya, as well as several from archaea. While more examples were found in bacteria and eukarya than archaea, this may just be a consequence of there being fewer archaea RNAs that have been studied with high resolution structural techniques. Of the 41 uniquely identified GU shifted G•U pairs, 39 were found in ribosomal RNAs. There presence in other classes of RNAs may be awaiting a more extensive 3D structural database for RNA. The shifted G•U wobbles were relatively rare compared to the standard wobbles. For instance, we identified 6,636 non-redundant standard G•U wobbles and 41 shifted G•U wobbles, or 0.61%. This rarity makes the shifted G•U of potential greater interest. For instance, it may be better to have a therapeutic target a rare motif rather than a common one, leading to less off-target binding and therefore greater specificity. Enhancing target specificity is a major issue in developing drug binding to RNA (53,54). Some of the shifted G•U motifs, such as Clusters 1 and 4 appear to be specific to bacteria, which might allow targeting without binding to the human host, either through directly targeting the shifted G•U or by using that G•U as a secondary binding site for two-domain drug binders (55).

While we identified 41 shifted G•U wobbles in the non-redundant dataset, there were many more such wobbles in the full dataset, 377 in total. As shown in Table 1, in each organism, we separated the G•U wobbles into two classes, shifted and standard, as judged by the hydrogen bonding scheme in Figure 4. It is notable that in the full dataset, 373 of these examples belonged to the shifted G•U wobble class, while only 4 were in the standard G•U wobble class. This suggests that the alternative protonation state(s) is dominant over the standard one, although other states or hybrid states outside of the shifted and standard G•U wobbles can also occur (Table 1). Other studies suggest that the anionic and enolic protonation states DNA and RNA have very low population (56,57), although those studies did not look at the fully shifted wobbles herein in which the G is resident in the major groove.

The nature of the alternative protonation state between anionic and enolic (Figure 1) is uncertain from the studies herein and will await NMR characterization. Unfortunately, reactivity of Us with DMS (Figure 9) does not inform on which of the alternative protonation states populate because the N3 of U is deprotonated, and therefore nucleophilic, in both anionic and enolic alternative states (Figure 1). It is possible that both anionic and enolic forms populate and that the former is favored at elevated pH.

It is notable that Clusters 1 and 3 both show one or more metal ions interacting with major groove G and U atoms (Figure 9), which suggests that the G•U may be in the anionic state in which the enolate of the U(O4) could contribute strongly to metal ion binding (Figure 1B). Future spectroscopic and theoretical studies will help resolve these questions. We also note that most metal ions were reported as Mg^2+^ ions, although a few as K^+^ ions. It is possible that the identity of the metal ion changes with the geometry of the G•U wobble, which will require future investigation.

Our analysis of the secondary structures of the 41 non-redundant shifted G•U wobbles revealed some prominent patterns. We formally divided the shifted wobbles into G•U and U•G subclasses but found striking symmetry between these. In the G•U subclass, there tended to be WCF pairs at the -1 sequence pair, while in the U•G subclass, there tended to be a WCF pair at the +1 sequence pair, albeit the G•U subclass had a predominance of GC WCF pairs while the U•G subclass had one of UA WCF pairs (Figure 6). The symmetry continued in that the +1 sequence pair in the G•U class tended to be unpaired, while the -1 sequence pair in the U•G class tended to be unpaired. Symmetry extended further to the -2 and +2 positions, which tended to be paired and unpaired in the G•U subclass respectively, but unpaired and paired the U•G subclass, respectively. The mirror image symmetry between the G•U and U•G subclasses, clear from comparison of the side-by-side panels B and C in Figure 6, is likely because one can rotate Figure 6A 180° about the axis coming out of the page to turn the G•U into a U•G and sequence pairs -1 and -2 into sequence pairs +1 and +2. Overall, one is left with a secondary structure motif in which the G•U sits atop a stem with a loop above it while the U•G sits below a stem with either unpaired or non-canonically paired nucleotides below it, making the symmetry imperfect.

Inspection of Figure 8, with its secondary structures, supports this model but with additional complexity. For instance, in the G•U subclass there can be a complex structure in the loop above that interacts with the G•U; for instance, cluster 3 contains a non-canonical trans-Watson-Crick/Hoogsteen AC base pair at sequence pair +1 that interacts with the G•U wobble. Likewise, in the U•G subclass there can be a complex structure in the stem below that interacts with the U•G; for instance, cluster 2 has a non-canonical cis-Watson-Crick/Watson-Crick AG base pair at sequence pair -1 that interacts with the U•G wobble. In some cases, the pairing neighboring the shifted wobble is relatively simple and strong; for example, clusters 1 and 3 share multiple GC base pairs above and below their respective U•G and G•U wobbles. But, in other cases, the pairing is more complex and longer; for example, in clusters 2 and 4 where cluster 2 has 4 non-canonical pairs below it and 9 mixed pairs above it, while cluster 4 has 6 mixed pairs below it.

These motifs raise the question as to what interactions are present in them to stabilize the alternative protonation states of the GU wobbles. Keeping with the theme that clusters 1 and 3 have similar secondary structures, they share key 3D structural features as well. For instance, there is a stabilizing metal ion in the major groove of each structure, although the wobble in cluster 1 has more sugar interactions and the wobble in cluster 3 has more minor groove interactions. Similarly, revisiting the theme that clusters 2 and 4 have similar secondary structures, they share essential 3D structural features too. For instance, there are extensive interactions with the sugars in each structure, with most of the G and U O2′ and O4′ atoms of the cluster members engaged in interactions, although the wobble in cluster 2 satisfies these with intrastrand interactions with neighboring nucleosides while the wobble in cluster 4 satisfies these primarily with amino acids from a ribosomal protein, albeit it does have one dominant intrastrand interaction with a neighboring nucleoside.

We asked what the dominant interactions in the shifted wobble might be that would direct the U towards the minor groove and the G towards the major groove. The structural clustering from the 3D structures reveals broad trends. In cluster 1, there were relatively few interactions in minor and sugar regions and only one major groove interaction, of the G with 5′-base. However, there were extensive metal ion interactions with both the G and the U. Thus, metal ions can play the dominant role in shifting the wobble. In cluster 2, there were extensive intrastrand major groove interactions of both the G and the U. In the minor groove, interactions were dominated by U, with a long-range interaction, while in the sugar region the interactions were dominated by the G, with both its O2′ and O4′. Thus, a plethora of interactions of both bases in the major, minor, and sugar region can shift the wobble. In cluster 3, the G dominates the major groove, again with extensive metal ion interactions. The minor groove has an intrastrand interaction between the G and the 3′-neighboring U, while there are no sugar interactions. Thus, once again metal ions interactions can play the dominant role in shifting the wobble, although here with just the G. Finally, in cluster 4, there were relatively few major groove interactions, two minor groove interactions, but a plethora of sugar interactions to the G and U. Unusually, these were largely with a ribosomal protein, albeit there is one dominant intrastrand interaction to the sugar of the U. This summary highlights that structural clustering can help identify diverse driving forces as being from preferential sequences, unique conformational states and atypical molecular interactions, all leading to the formation of a shifted G•U wobble. Finally, we note that the cheminformatics approach developed herein can be applied to other non-WCF base pairs to identify additional novel protonation states of RNA and is applicable to DNA and proteins as well.

## Acknowledgements

The authors would like to thank Dr. Andrey Krasilnikov for advice on the analysis of RNA structures. We also thank Kobie Kirven for advice on computational and statistical analysis and Dr. Mrityunjay Gupta for helpful comments on the manuscript.

## Funding

This research was supported by the National Institutes of Health (NIH) grant R35GM127064 and the National Aeronautics and Space Administration (NASA) grant 80NSSC22K0553.

## Supplementary data

Will be made available upon peer review of the manuscript.

## Notes

### Competing Interest Statement

The authors have declared no competing interest.

